# Direct observation of the setular web that fuses thoracopodal setae of a calanoid copepod into a collapsible fan

**DOI:** 10.1101/808196

**Authors:** George von Dassow, Richard B. Emlet

## Abstract

Copepods are numerically dominant planktonic grazers throughout the waters of Earth, preyed upon in turn by a wide diversity of pelagic animals (1,2). Their feeding and swimming performance thus has global importance to aquatic food webs and oceanic carbon flux. These crustaceans swim and feed using cuticle-covered, segmented, muscular appendages whose reach is extended greatly by setae, extracellular chitinous extensions with diverse structure and function (3). Plumose setae, with subsidiary setules arranged like barbs on a feather, have well-documented roles in generating feeding and swimming currents (4,5). Recent work showed that plumose setae of barnacle cyprid thoracopods are permanently linked by setules into a single fan that opens and closes as one sheet during high-speed swimming (6). Intersetular linkage across cyprid thoracopods may greatly decrease leakage between extended setae, ensure even spread of setae within the fan, and promote ordered collapse of the fan to avoid entanglement of adjacent appendages. Here we demonstrate similar setular webbing amongst thoracopod setae in the calanoid copepod *Acartia* sp. High-speed video directly documents the existence of such links, and reveals that individuals experience apparently-irreparable degradation of the setal array due to de-linkage, with likely consequences for swimming performance.

---

We caught calanoid copepods and copepodites in plankton samples from the Charleston marina, Oregon, USA (43° 20.682’N, 124° 19.236’W). *Acartia* sp. were most prevalent, and were first concentrated by attraction to light, then selected individually after anaesthetization in MgCl_2_ or tricaine. To record escape swimming, one or several animals were introduced into 0.3-0.5 ml clean natural seawater in a coverslip-bottomed 35 mm Petri dish (sometimes seeded with *Nannochloropsis* as a neutral, non-motile, non-aggregating particulate tracer), then covered with another coverslip supported by large clay feet, and viewed on an inverted microscope. Animals were allowed at least two body lengths of vertical distance in such chambers (vertical depth: 2.5-4 mm). Low-magnification recordings revealed the use of appendages during escape swimming (Fig. 1 and video 1): each episode begins with a posterior sweep of the antennules, accompanied or immediately followed within ~10 ms by a metachronal strokes of four pairs of setose thoracopods. During the power stroke, the major setae of each thoracopod pair spread into a fan with nearly-even spacing between radially-projecting setae, spanning as much as a half-circle at full extent (Fig. 1, insets 1 and 2). As they sweep posteriorly, appendage pairs fold medially (Fig. 1, inset 3), then setae collapse together (Fig. 1, inset 4). During recovery, all thoracopods move together, with setae bundled tightly into brushes (Fig. 1, inset 5). The entire sequence of power and recovery strokes of all four thoracopod pairs is completed on the order of 10 ms. As described previously (7), episodes often consist of multiple power-recovery sequences; termination is accompanied by restoration of antennules to laterally-extended posture; and in most escape swim episodes, the urosome commences repeated dorsoventral flexure, with caudal rami outspread, after the first set of thoracopod strokes.

**Figure 1.**
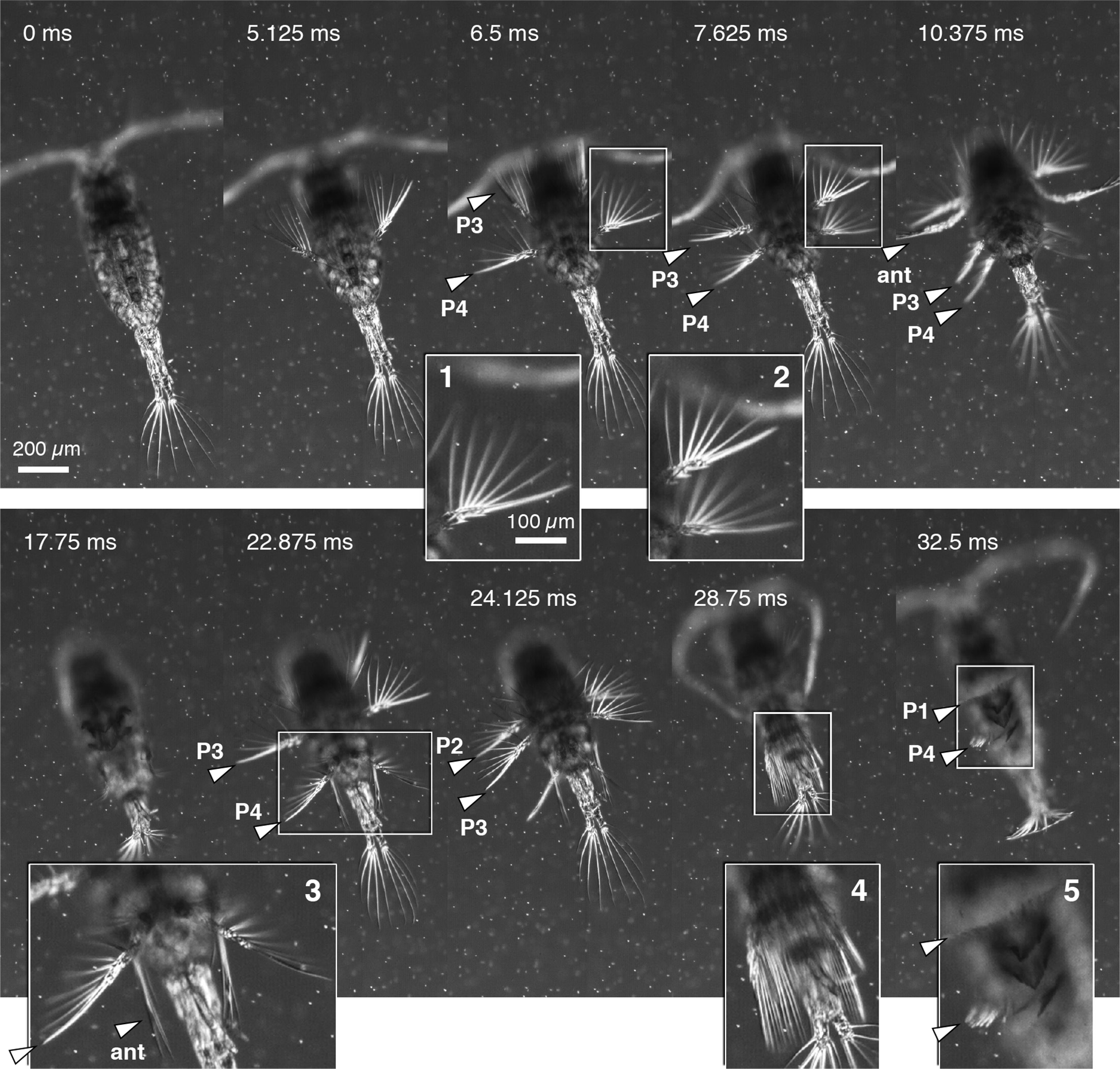
General characteristics of *Acartia*’s escape swim. Frames from a low magnification DIC sequence (Video 1) of a swim episode involving two successive power-recovery cycles within ~30 ms. Medium seeded with *Nannochloropsis*. Insets: 1=exopodite of P4 near full extent; 2=exopodites of P3 and P4 near full extent; 3=P4 folding medially, fan still extended (ant=antenna); 4=setae of all appendages nearly bundled; 5=bundled setae (P4 not quite fully bundled, P1 still slightly extended).

To detect setular webbing in live, intact animals requires exceptionally high-speed video because, while anaesthetized animals do hold their thoracopods extended, they do so in an unnatural posture in which the setal fan remains bundled. Natural thoracopod movement in *Acartia* is so fast that it required exposures on the order of 100,000th of a second to limit motion blur, which otherwise obscured the fine setular mesh between setae. This short exposure time precluded use of contrast enhancement techniques such as DIC, hence we relied on Köhler illumination with low aperture to increase contrast and depth of field. Even at modest magnification, it is apparent that setules form a mesh between adjacent setae, and that deviations from regular spacing within the fan reflect defects in this setular webbing (Fig. 2A, inset 1 vs. 2, and video 2).

**Figure 2.**
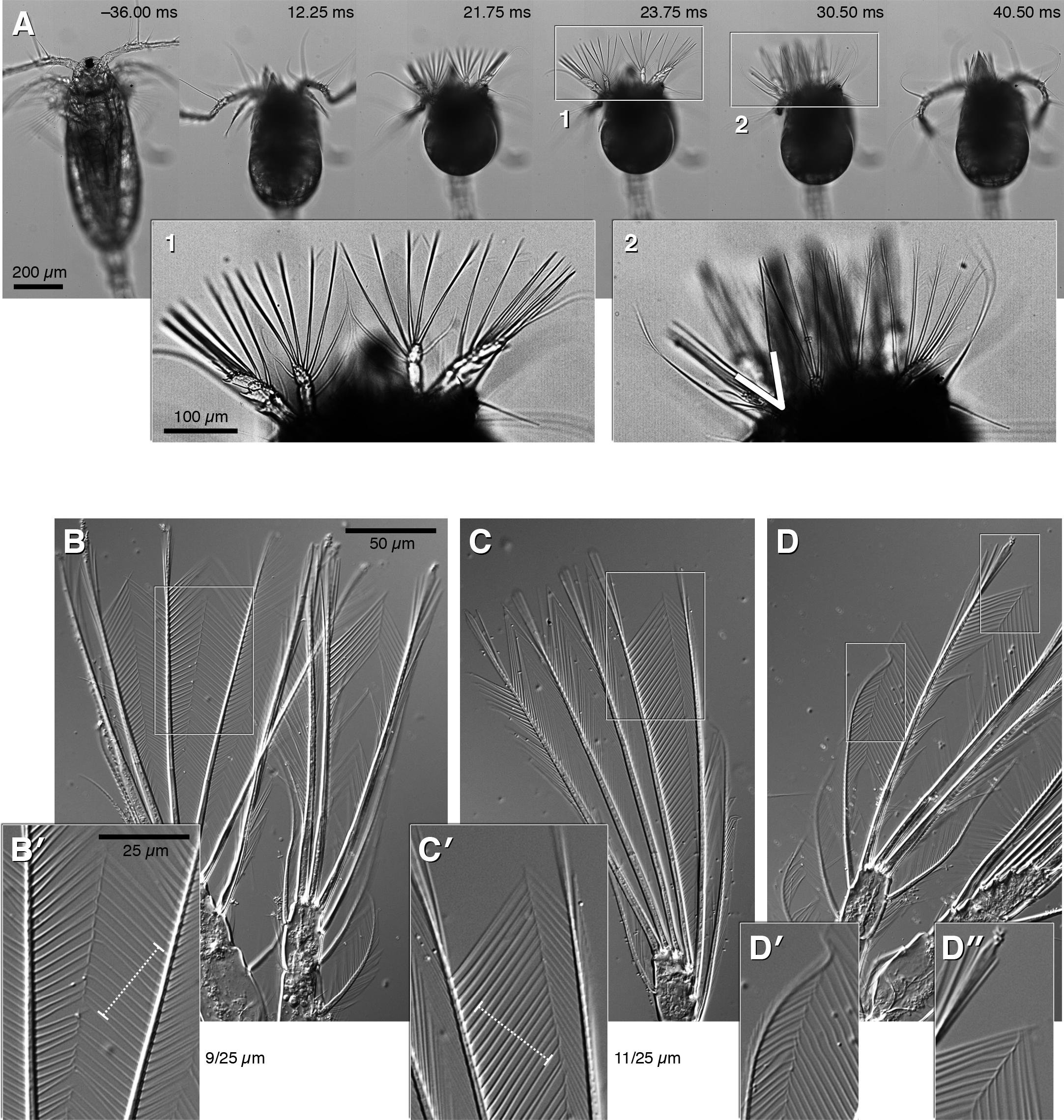
Setules linked across the setal fan. A) Frames from brightfield sequence in nearly full axial view (corresponds to Video 2). Inset 1 shows P2 with near-uniform spread of medial setae, mesh visible between them; Inset 2 shows P1 with some left-side setae spread too far apart (white V) and without visible mesh between them. B–D) Dissected appendages spread under coverslip, viewed by DIC. B) P1, with exopodite setae spread (inset B’); C) endopodite of P2 or P3, major setae spread (inset C’); D) endopodite of P2 or P3 highlighting medial setae (inset D’) and between adjacent major setae (inset D”). Dashed “calipers” are 25 μm, with adjacent counts of spanned gaps.

To confirm that the mesh represents physical linkage amongst setules, thoracopods were removed from anesthetized animals at or near the base with fine forceps, then spread on a slide in a small drop of water by application of a coverslip, as previously done with barnacle cyprids (6). In such preps, we could demonstrate linkage amongst all thoracopod setae in *Acartia* (Fig. 2B-D), although compared to barnacle cyprids the links were frail and easily broken by the tension applied to spread the array. Indeed, most links broke as we watched, with the consequence that most de-linked setules snapped flush to the setal shaft, indicating an elastic region at the base of each setule. In spread preps of dissected appendages we could not detect setular linkage in any of the head appendages, nor conclusively in the caudal rami (although some video sequences suggest that some linkage may exist amongst caudal setae). Notably, caudal rami and maxillary setae retained the vaned form when spread; that is, we did not detect any elastic joint between setule and setal shaft, unlike in the thoracopods.

Escape sequences captured at 4-8,000 fps with 1/100,000 sec exposure time clearly showed both the existence of setular linkage between all thoracopod setae, and also showed that these links flex as the array opens and closes (Fig. 3 and video 3). Especially striking are the links between the smaller setae on the medial side of the endopodites. These stitch together the two appendages of a pair along the midline, making one single fan out of the entire array (Fig. 3, inset 2). This is directly analogous to the stout setules that link appendage pairs in the barnacle cyprid, but instead relies on many fine links instead of a few stout ones. That these medial setae are linked is shown unambiguously by their deep flexure as the fan extends to full width (video 3). Near full extension there are 10-12 setules per 25 microns (Fig. 3), measured perpendicular to the setules (10.9±1.3 setules/25 μm, 44 measurements from 16 live recordings across all appendage pairs and zones within the setal fan). Assuming at least 0.5 micron for the setule itself, this means the largest fluid aperture within an intact thoracopod fan is likely no more than 2 microns. These apertures are erected across the fan in ~1 ms, remain open for a few milliseconds, and become occluded as setules collapse onto parent setae in the next 1 ms in preparation for the recovery stroke.

**Figure 3.**
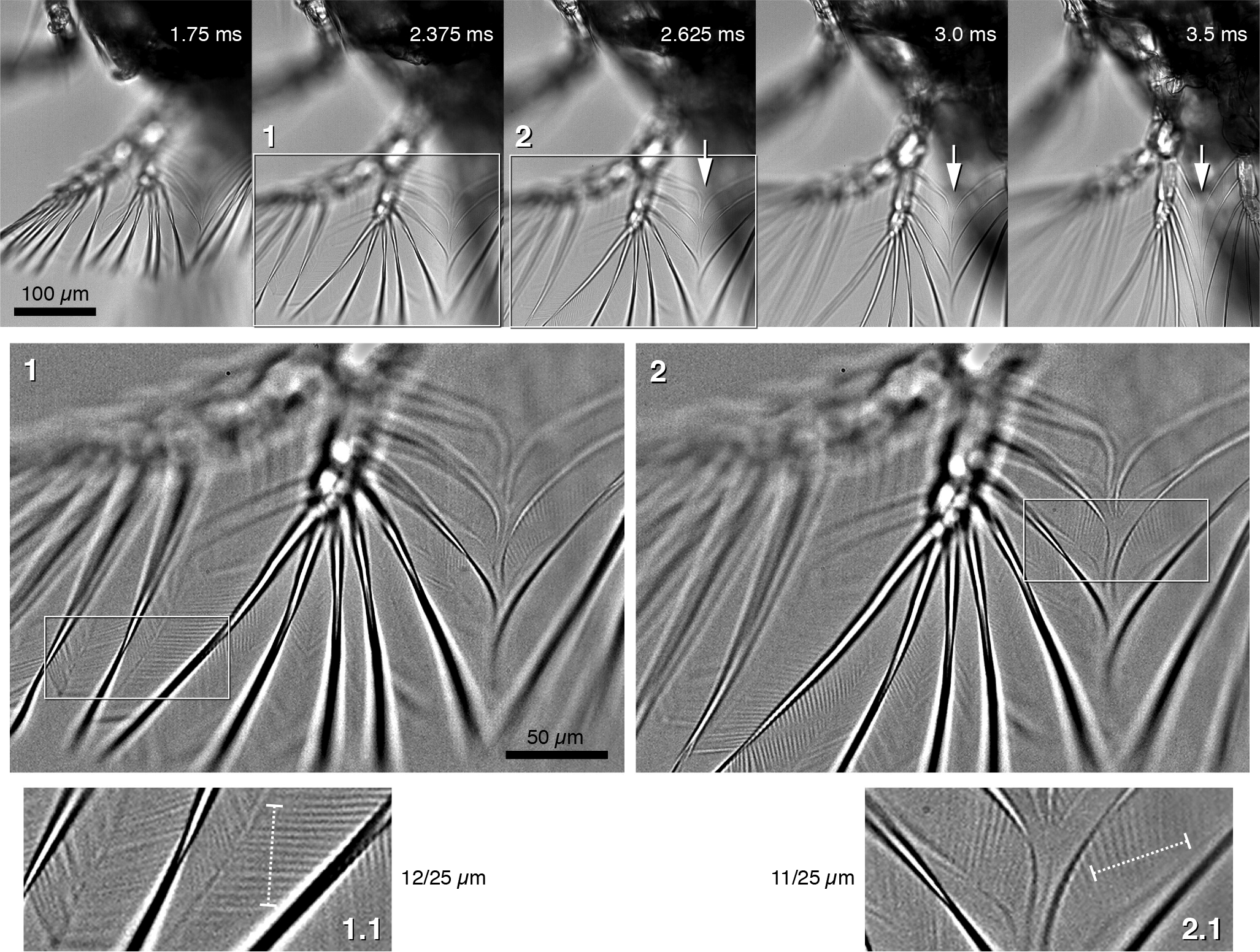
Setular linkage in live recordings. Frames show appendage pair P2 in an individual with nearly intact setular mesh in approximately axial view (see Video 3), from partially extended (first frame) through maximal spread (third frame), to folding medially (fifth frame). Insets were high-pass filtered and contrast-enhanced to highlight setules. These frames capture the dramatic flexure of the medial setae borne by the shank of the endopodite as the fan extends, then their straightening as it folds (arrows). Flexure must reflect linkage by setules, without which these would interdigitate. Furthermore, if similar flexure were observed in animals where the mesh cannot be directly observed, it would be indirect evidence of setular linkage. Inset 1 captures the zone between exopodite and endopodite as they pass through focus; inset 2 captures the medial fusion between the setae of left and right appendages. 2x blowups (1.1, 2.1) illustrate the measurement of spacing in the setular mesh: counting gaps in a 25 μm distance, perpendicular to setules, with array near maximal spread and parallel to focal plane.

Most animals exhibited extensive linkage across the entire thoracopod array on the day of collection. However, defects became more prominent once animals had spent a day (or more) confined in a sample beaker. Degradation of setal fans was evident in most cases, and included broken setal tips, stretches of unlinked setae within the fan (Fig. 4 and video 4), and cases in which adjacent setae had become entirely unzipped from one another. The latter category is visible even at low magnification, because it results in an excessive angle between the unlinked setae and failure to spread in the next one or more setae beyond (Fig. 2A, inset 2). These cases also show that the fanning-out is in part due to the spread of exopodite from endopodite, as the corresponding sections of the fan are isolated from one another with respect to defects.

**Figure 4.**
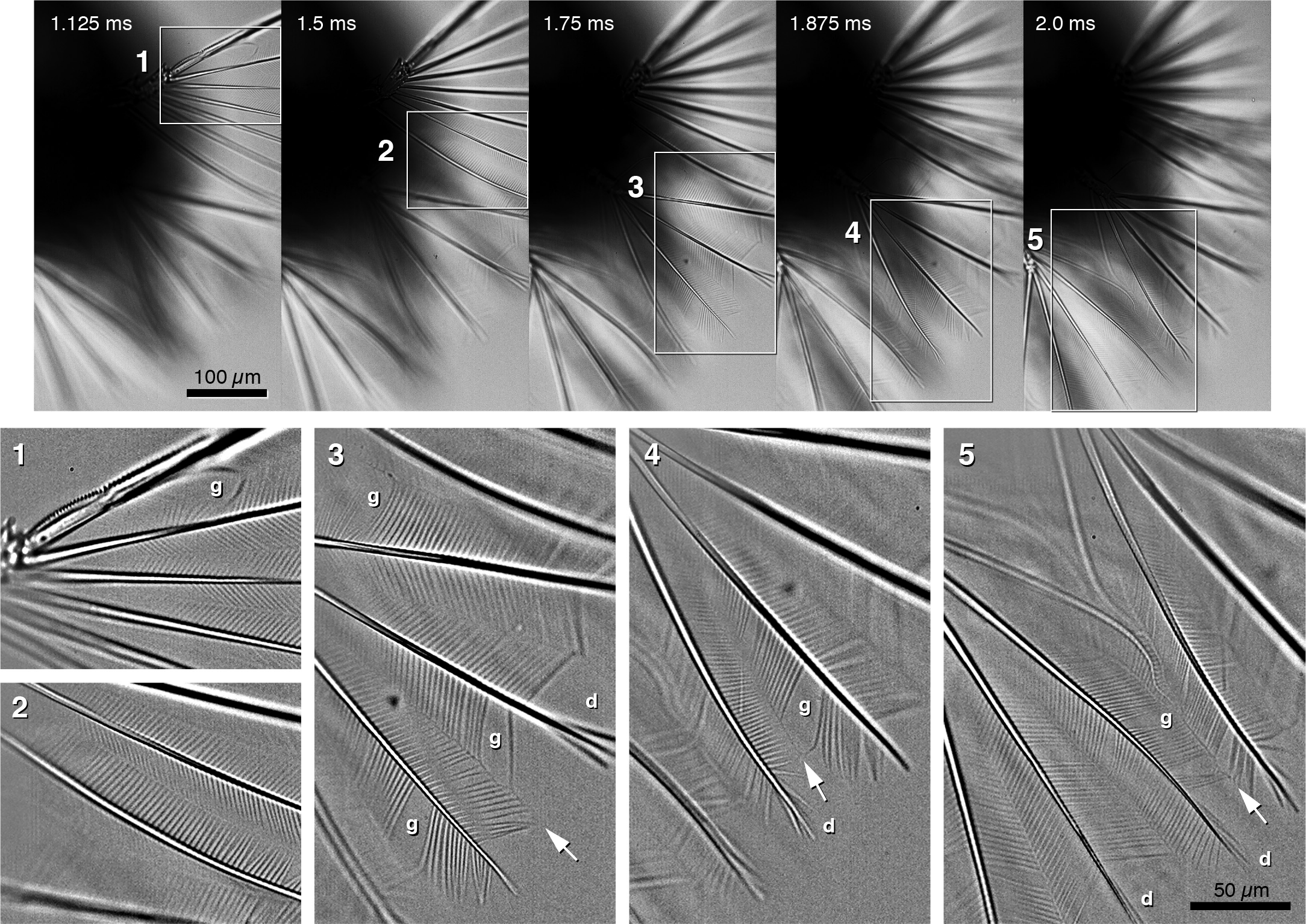
Degradation of the setular mesh. Frames show appendage pair P4 in an individual with small gaps (g) and distal zones of de-linkage (d) in the setular mesh (see Video 4). View otherwise as in Fig. 3; insets are high-pass filtered and contrast-enhanced. Curiously, in some gaps a visible thread remains where setules from adjacent setae would have met (arrows, insets 3–5). We cannot detect barbs or hooked ends on free setules either in video frames or when are spread under coverslips; yet when linked, setule tips appear to flex when strained (see Fig. 2B’). Hence we suspect this remnant thread represents some kind of cement, or broken-off tips of setules still embedded in such cement, that has been somehow applied to unite setules in register.

The existence of setular linkage could also be demonstrated in tricaine-anesthetized animals, which hold their thoracopods in an unnaturally-extended posture with bundled setae. Plucking the setae with a fine glass needle extended one or more sections of the fan, spreading linked setules between them (Fig. 5); tellingly, when multiple sections could be spread, the spacing between them was approximately even (Fig. 5A), which we interpret to be due to the elasticity of the setular mesh. Conversely, when sections were stretched, setular links broke (Fig. 5B). As their links failed, setules swung distally to lie alongside the parent seta, which we interpret as an elastic joint between setule and parent seta that tends to close the fan. Once the entire setule set became unlinked, the adjacent seta swung medially, implying elasticity at the setal base that also tends to close the fan. Similarly probing setae of the caudal rami resulted in mere separation of the unlinked plumes, and showed that the setules thereon return to vaned orientation when strummed by the needle.

**Figure 5.**
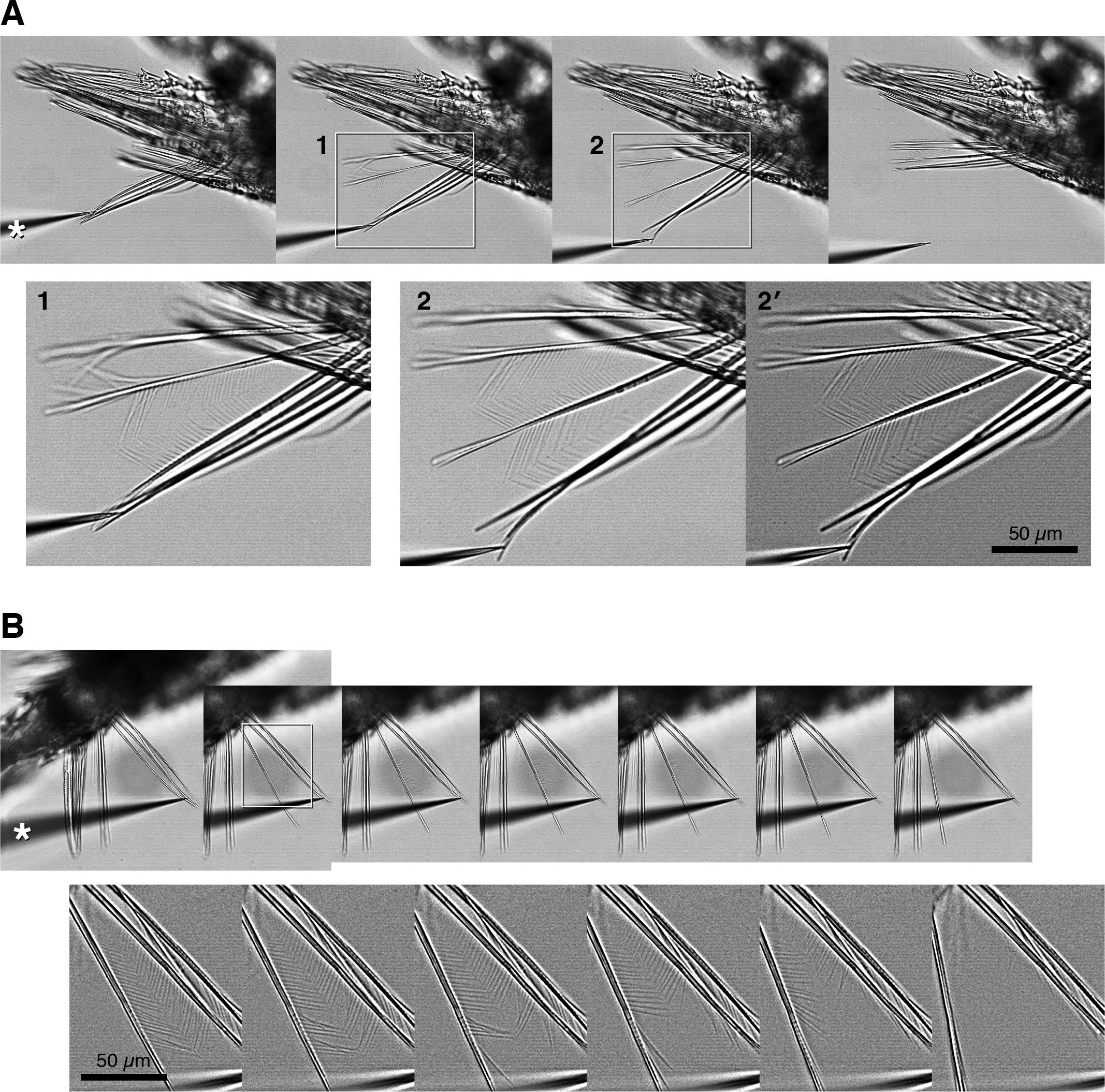
Micromanipulation demonstrates consequences of linkage. Tricaine-doped animals held in place by vacuum grease were probed with a glass needle (asterisks); stage control was used to move the animal abruptly relative to the needle. A) Setae from endopodite of P2 or P3 first pull free from the array, then partially spread as each seta slips from under the needle. As first one, then another seta is freed, they are restrained by the setular mesh. Insets 1 and 2 are unprocessed but contrast enhanced; 2’ is high-pass filtered. B) A similar endopodite strained to the point that setules unzip. High-pass filtered blowups of boxed region show progressive release; as setules become unlinked, they swing distally to lie along the parent seta.

Finally, seeding the medium with *Nannochloropsis* (a non-motile alga in the few-micron size range) illustrates that these copepods, using the webbed thoracopod setal fan, reach speeds that exceed the viscous-dominated regime that nearly universally governs movement of small pelagic animals. Inertial motion of suspended particles was readily apparent during escape swimming, at least after the thoracopods had conducted a power stroke (video 5). In a few cases, however, we detected particles passing through the fan via large defects, i.e. gaps between unzipped setae (video 6). These instances imply that progressive degradation of the setular web may occur between molts and likely compromises swimming performance as leakage increases.

Previous observers of copepod swimming may not have fully recognized the factors by which power and recovery strokes of the thoracopods differ. Two well-recognized factors, visible even in low-magnification high-speed video, are 1) the metachrony of the power stroke versus the synchrony with which collapsed appendages return to starting position (8,9,10), and 2) the buckling of appendage joints such that the recovery stroke undercuts the power stroke’s sweep (5). Less well-documented is 3) the lateral spread of thoracopod setae into a fan (but see 10,11,12). If the newly-discovered setular mesh between the setae effectively weaves them into a solid fan, the difference in area – at least several-fold – multiplies the effect of metachrony and stroke asymmetry. The degree to which the setal fan enhances propulsion depends on the leakiness of the fan (13), and whether motion is viscous-dominated or inertial. In the viscous-dominated regime, spreading thoracopod setae from a bundle into an array of isolated rods increases drag, assuming that setae are stiff enough to remain erect during the power stroke, and hence moves more water per stroke. In contrast, in the inertial regime, isolated rods may not propel much more fluid than the bundle. Hence, the potential role of the setular webbing: by greatly reducing or eliminating leakage between setae, the setules convert what would be a rake into a paddle.

*Acartia*’s webbed thoracopod fan is strikingly similar to traits recently discovered in barnacle cyprids (6). Despite differences in anatomy, the apparent functions of setular linkage in swimming are the same. This begs the question of how widespread a trait it might be. Lamont and Emlet (6) described it in all barnacle cyprids (including acrothoracicans, rhizocephalans, and thoracicans; no information is available for basal thecostracans), but not in nauplii. We have not detected linkage in any of several decapod zoeas, other small pelagic decapods, or harpactacoid copepods. Even in large *Acartia*, the setular webbing is on the threshold of visibility. In calanoids other than *Acartia* the setules were difficult to image in live animals either because they were too fine to show up at the relatively low resolution, high light, and short exposure required to have any chance of catching an untethered copepod in the field of view, or because the animals move too fast for our equipment to overcome motion blur. From the flexure of the medial setae during the power stroke, however, we believe that setules likely link thoracopod setae in other calanoids as well. We also tentatively detected linkage in high-speed recordings of the cyclopoid *Oithona similis*, but could not confirm it with amputated appendage preps.

We hypothesize that setular linkage is an essential adaptation enabling these animals to achieve the remarkably high speed swimming that previous authors called “escape from viscosity” (8). If setular linkage optimizes escape swimming, this raises an important ecological question. It is presently difficult to imagine how these animals might effectively maintain setular linkage between molts. If they cannot, then – leaving aside the mystery of how such linkage might arise during cuticle formation in the first place – it seems likely that defects large and small, ranging from a few broken setules to entire disconnection of adjacent setae or even breakage of multiple setae, must accumulate after molting. This implies in turn that escape swim performance must suffer a concomitant degradation, imposing as-yet-unexplored costs of activity and pressures on the intermolt duration for copepods or the time to settlement for cyprids.

## Supplemental Videos

Videos 1–4: see legends to Figures 1, 2A, 3, and 4.

Video 5: Low magnification sequences of multi-stroke swims in sewater seeded with *Nannochloropsis*; first segment, frontal view, second segment in side view. Each segment is shown unprocessed, then overlaid with particle trails, then particle trails alone. Trails were created by Fourier band-pass filtering images, then coding the finest band in red, the next-finest in green, and the coarsest in blue, such that particles appear reddish when closest to focus, and blueish when far from focus. Trails are 24-frame maximum projections.

Video 6: Low magnification DIC sequence of an animal with significant setular degradation, swimming in seawater seeded with *Nannochloropsis*. Second loop slows play and highlights, on Fourier-filtered insets, particles that appear to slip between setae due to mesh defects. Three are highlighted per stroke, but examination frame-by-frame shows more that slip through.

